# GNS561, a New Autophagy Inhibitor Active against Cancer Stem Cells in Hepatocellular Carcinoma and Hepatic Metastasis from Colorectal Cancer

**DOI:** 10.1101/2020.12.21.423741

**Authors:** Sonia Brun, Jean-Marc Pascussi, Elena Patricia Gifu, Eloïne Bestion, Zuzana Macek-Jilkova, Guanxiong Wang, Firas Bassissi, Soraya Mezouar, Jérôme Courcambeck, Philippe Merle, Thomas Decaens, Julie Pannequin, Philippe Halfon, Claude Caron de Fromentel

## Abstract

Patients with advanced hepatocellular carcinoma (HCC) or metastatic colorectal cancer (mCRC) have a very poor prognosis due to the lack of efficient treatments. As observed in several other tumors, the effectiveness of treatments is mainly hampered by the presence of a highly tumorigenic sub-population of cancer cells called cancer stem cells (CSCs). Indeed, CSCs are resistant to chemotherapy and radiotherapy and have the ability to regenerate the tumor bulk. Hence, innovative drugs that are efficient against both bulk tumor cells and CSCs would likely improve cancer treatment. In this study, we demonstrated that GNS561, a new autophagy inhibitor that induces lysosomal cell death, showed significant activity against not only the whole tumor population but also a sub-population displaying CSC features (high ALDH activity and tumorsphere formation ability) in HCC and in liver mCRC cell lines. These results were confirmed *in vivo* in HCC from a DEN-induced cirrhotic rat model in which GNS561 decreased tumor growth and reduced the frequency of CSCs (CD90^+^CD45^-^). Accordingly, GNS561, which was in a global phase 1b clinical trial in liver cancers that was recently successful, offers great promise for cancer therapy by exterminating both the tumor bulk and the CSC sub-population.

## Introduction

Patients with advanced hepatocellular carcinoma (HCC) or metastatic colorectal cancer (mCRC) have a very poor prognosis [1]. They suffer from a lack of efficient therapy for HCC or treatment failures for CRC, both resulting in low survival rates. Since the late 1990s, a sub-population of poorly differentiated cancer cells, known as cancer stem cells (CSCs), have raised substantial interest, as they are believed to play a crucial role in cancer initiation, propagation, relapse and metastasis [2–5]. According to the stem cell paradigm, CSCs are able to both self-renew and give rise to progenitors thus to generate the bulk of a tumor ad infinitum [6]. This model has been validated for several tumor types, including HCC and CRC, whereas other ones, such as melanoma [7], do not exactly follow this pattern [8]. It has also been shown that CSCs are resistant to various types of stress, including those generated by treatments such as radiotherapy or chemotherapy [9,10]. Recently, it was shown that apoptosis was induced in the majority of HCC tumor cells following treatment with sorafenib, a standard of care, whereas a subpopulation of CSCs remained and led to tumor metastases [11,12]. In the same line, colorectal CSCs are not sensitive to radiotherapy or chemotherapy with oxaliplatin, irinotecan or 5-FU, leading to tumor metastasis and recurrence after chemotherapy [13]. Thus, innovative drugs that simultaneously exterminate both bulk tumor cells and CSC subpopulation can improve cancer treatment and result in long-term tumor remission [4,8].

We previously reported that GNS561, a new lysosomotropic molecule with high hepatic tropism, was efficient against intrahepatic cholangiocarcinoma and HCC [14,15]. Based on robust preclinical data, the potential use of GNS561 against primary and secondary liver cancers was evaluated in a global phase 1b clinical trial (NCT03316222), which was recently successful [16]. Regarding its mode of action, GNS561 was shown to suppress cancer cell growth *via* the induction of apoptosis resulting from inhibition of palmitoyl-protein thioesterase 1 (PPT1) activity, lysosomal unbound Zn^2+^ accumulation, impairment of cathepsin activity, blockage of autophagic flux, altered localization of mTOR, lysosomal membrane permeabilization and caspase activation [14].

The inhibition of autophagy induced by GNS561 suggests that this molecule could be effective in killing CSCs. Indeed, autophagy, as a catabolic and prosurvival pathway, appears to be critical for the survival of cancer cells and especially for the maintenance, plasticity, and chemoresistance of CSCs and their adaptation to tumor microenvironment changes (TME) [3,5,17,18]. An increase in autophagy levels as a source of nutrient replenishment was reported in cancer cells and CSCs in response to increased metabolic demands triggered by environmental stress signals, such as hypoxia, starvation, metabolic and oxidative stresses and therapy [19,20]. Therefore, autophagy inhibition might be a useful strategy for targeting and killing both cancer cells and CSCs [3,21–23].

In this study, we investigated the effect of GNS561 on CSCs. We first showed that *in vitro*, GNS561 induced a dramatic increase of CSC death in both HCC and liver mCRC cell lines. We then confirmed its efficacy on CSCs *in vivo* in an HCC-induced cirrhotic rat model.

## Materials and Methods

Details of the materials and methods are provided in the Supplementary Methods.

### Cell line culture

Hep3B cells were maintained in EMEM (ATCC) supplemented with 10% heat-inactivated fetal bovine serum (FBS) (Invitrogen) and 1% penicillin/streptomycin (Gibco). Huh7 cells were cultured in DMEM-F12 medium (Gibco) supplemented with 10% heat-inactivated FBS (Invitrogen), 1% nonessential amino acids (Gibco), 1% penicillin/streptomycin (Gibco), 1% GlutaMAX (Invitrogen) and 1 mM sodium pyruvate (Gibco). Liver mCRC patient-derived cell lines (CPP19, 30, 36 and CPP45) [24] were maintained in DMEM (Gibco) with 10% FBS. All cells were cultured in a humidified atmosphere at 37 °C and 5% CO_2_.

### Diethylnitrosamine-induced cirrhotic immunocompetent rat model of HCC

Rats with HCC were treated with vehicle, GNS561 or sorafenib, by oral gavage for a period of six weeks. The animals were checked daily for clinical signs, effects of tumor growth and any other abnormal effects. All rats received humane care in accordance with the Guidelines on the Humane Treatment of Laboratory Animals (Directive 2010/63/EU), and experiments were approved by the animal Ethics Committee: GIN Ethics Committee No. 004.

### Statistical analysis

Statistical analyses were performed using Prism 8.4.3 software (GraphPad Software, Inc.). For datasets with a normal distribution, multiple comparisons were performed using one-way ANOVA with Dunnett’s post hoc analysis. For datasets without normal distribution, multiple comparisons were performed using the non-parametric Kruskal-Wallis test. Data are presented as the mean values ± standard error mean (SEM) of three independent experiments unless stated otherwise. Statistical significance is indicated by one (P-value < 0.05) or two (P-value < 0.01) asterisks on the figures.

## Results

### GNS561 is efficient against the whole tumor

We first investigated the activity of GNS561 in the whole population of two HCC cell lines and four liver mCRC cell lines. GNS561 decreased viability in a dose-dependent manner, with very similar maximal inhibitory concentration (IC_50_) values for all the tested cell lines and < 3 μM (Table 1).

**Table 1.** *In vitro* activity (IC_50_) of GNS561 on two HCC (Hep3B and Huh7) and four liver mCRC cell lines (CPP19, CPP30, CPP36 and CPP45) after 72 h of treatment with GNS561.

| | Name | $IC_{50}$ ($\mu M$) | 95% confidence interval |
| --- | --- | --- | --- |
| HCC cell lines | Hep3B | 2.33 | 2.358 to 2.535 |
|  | Huh7 | 1.48 | 1.238 to 1.699 |
| Liver mCRC cell lines | CPP19 | 1.88 | 1.424 to 2.487 |
|  | CPP30 | 1.22 | 0.9210 to 1.624 |
|  | CPP36 | 1.45 | 0.6990 to 3.025 |
|  | CPP45 | 1.12 | 0.9555 to 1.318 |

### GNS561 is also active against a subpopulation displaying CSC features

Aldehyde dehydrogenase (ALDH) is known to be a marker of CSCs in numerous solid cancers, including CRC [25,26]. To evaluate the sensitivity to GNS561 of the CSC/cancer progenitor cell-enriched subpopulations of liver mCRC cell lines, the percentage of ALDH^bright^ cells was quantified by flow cytometry after 72 h of incubation with GNS561. As shown in Fig. 1A, GNS561 significantly decreased the percentage of ALDH^bright^ cells in a dose-dependent manner in all three mCRC cell lines.

**Figure 1.**
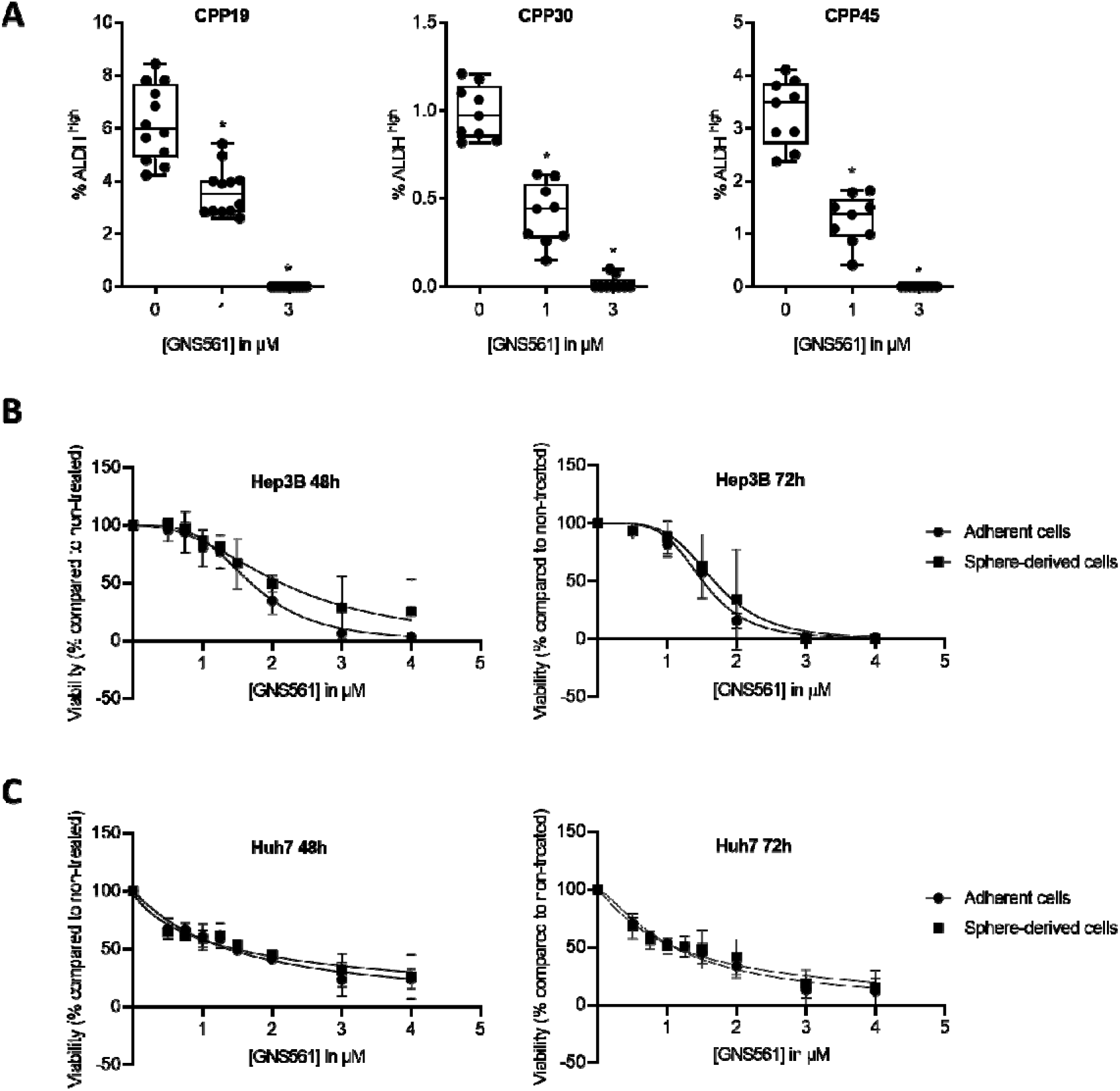
GNS561 is efficient against subpopulation displaying CSC features. (A) Box and whisker representation (min to max) of the percentage of ALDH^bright^ cells after 72 h of treatment with GNS561 in three liver mCRC cell lines with the indicated concentrations. For comparison with the control, one-way ANOVA with Dunnett’s post hoc analysis was performed. *, P < 0.05. Antitumor activity of GNS561 on whole tumor (circle) or CSC-enriched populations (square) in Hep3B (B) and Huh7 (C) HCC cell lines after 48 or 72 h of treatment with GNS561. The curves represent the mean of two independent experiments in triplicate.

Unlike CRC cells, ALDH-overexpressing HCC cells seem to exhibit a differentiated rather than a CSC phenotype [27]. Moreover, there is a lack of consensus biomarkers for CSCs in HCC [23,28]. Consequently, the CSC subpopulation is commonly identified in HCC based on functional criteria such as their capacity to efflux Hoechst 33342 dye (side population) [29] or to form tumorspheres [30,31], as only these cells have the ability to proliferate in non-adherent conditions [32,33]. Tumorsphere formation assay was used to isolate the CSC subpopulation of HCC cell lines. The significant overexpression of the stemness-associated factors *SOX2* and *NANOG* in Hep3B tumorspheres compared to the whole cell population confirmed their immature phenotype (Fig. S1). To avoid chemoresistance because cells within the interior of tumorspheres are protected from drug penetration by neighboring cells on the periphery of the sphere, we compared the impact of GNS561 between the whole and CSC-enriched populations of Hep3B and HuH7 cell lines in monolayer cultures. GNS561 had the same effect on the viability of the two populations from Hep3B (Fig.1B) or Huh7 (Fig.1C) cells treated in monolayer cultures for 48 or 72 h. Since cells with an immature phenotype have high plasticity when cultured *in vitro*, we hypothesized that they could have derived after 48 or 72 h of monolayer culture and spontaneously restored the initial heterogeneous population [6]. Thus, we repeated the experiment in Hep3B cells and assessed the antitumor activity of GNS561 at shorter times (6, 8 and 24 h) and at higher doses (0.5 to 32 μM). Similarly, GNS561 was as efficient against the bulk population as the CSC-enriched population (Fig. S2).

### GNS561 blocks tumorsphere formation

Furthermore, the ability of GNS561 to impair tumorsphere formation was investigated in both liver mCRC and HCC cell lines. GNS561 (0.3 or 3 μM) was added after seeding a low number of liver mCRC cells in CSC medium. Ten days later, tumorspheres > 50 μm were counted. As shown in Fig. 2A and 2B, GNS561 induced significant dose-dependent decreases in tumorspheres in all three cell lines, with complete suppression of tumorspheres at the highest dose (3 μM). Similar results were obtained with the Hep3B HCC cell line at 96 h (Fig. 2A and B) and 11 days (data not shown) of GNS561 treatment. As illustrated in Fig. 2, GNS561 completely inhibited tumorsphere formation, even at low concentration (0.8 μM corresponding to 1/3 IC_50_ in the adherent whole population). These results showed that GNS561 has an antitumor effect not only on the bulk, but also on the CSC-enriched subpopulation. In order to further enrich the CSC population, secondary spheres were generated from primary Hep3B spheres and treated with the same doses of GNS561. A slight difference in sensitivity was observed between primary and secondary spheres, with a complete inhibition of sphere formation reached at 0.8 and 1.6 μM, respectively (Fig 2B and Fig.S3). These results confirm the significant activity of GNS561 on CSCs. In contrast, sorafenib had only a mild effect on the CSC-enriched subpopulation since it failed to completely abolish tumorsphere formation whether from adherent cells or primary spheres (Fig. S3 and S4).

**Figure 2.**
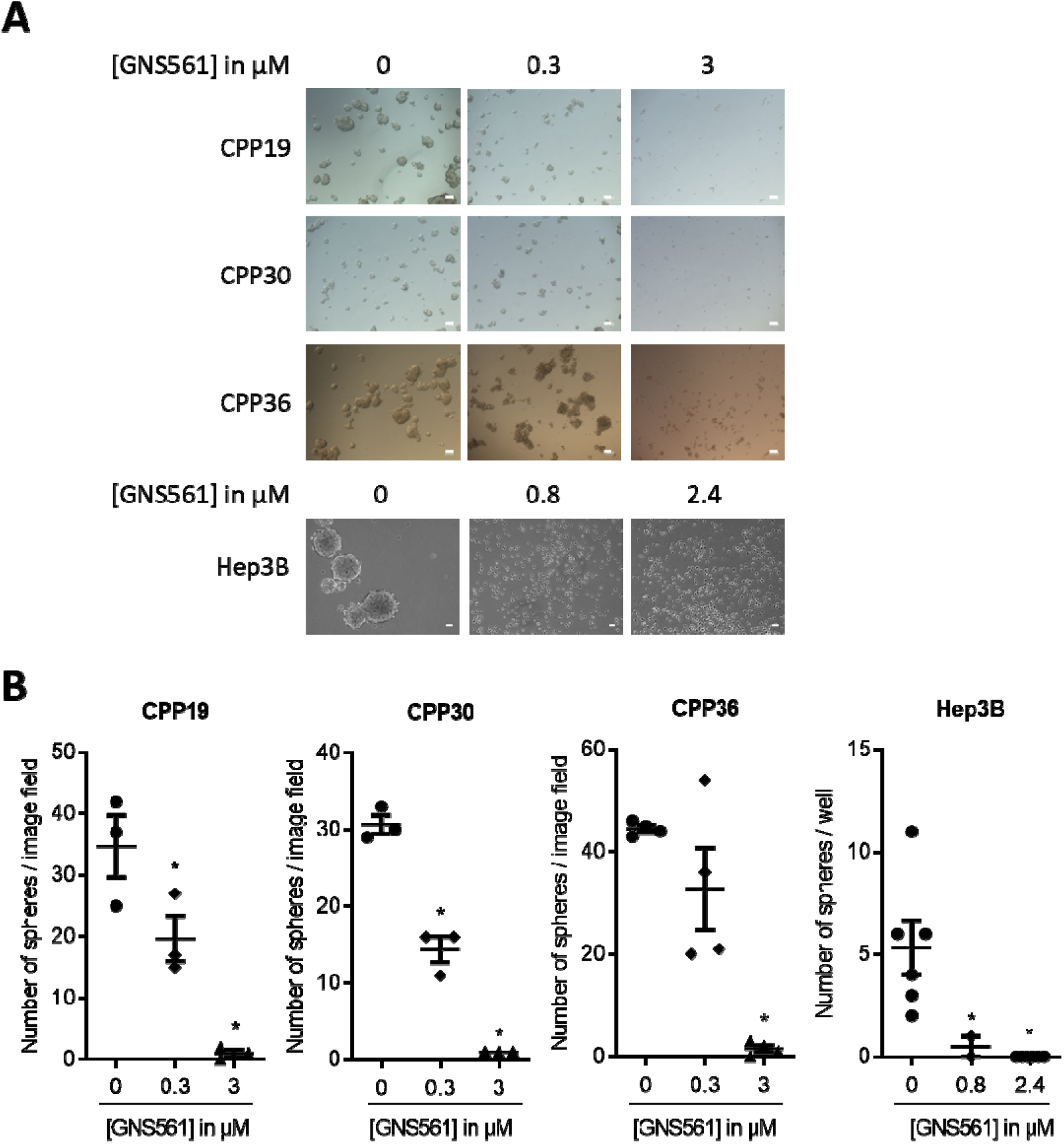
GNS561 alters the self-renewal of liver mCRC and HCC cell lines. (A) Tumorsphere forming efficiency of three liver mCRC (CPP19, CPP30 and CPP36) and one HCC (Hep3B) cell lines treated with the indicated GNS561 concentrations. Scale bars represent 50 μm. (B) Number of spheres > 50 μm. For comparison with the control, statistical significance was determined using one-way ANOVA and Dunnett’s post hoc test. *, P < 0.05.

### GNS561 decreases tumor growth and CSCs in a diethylnitrosamine-induced cirrhotic rat model of HCC

As demonstrating the antitumor activity of a drug on CSCs by tumorsphere formation *in vitro* could reflect its *in vivo* anti-tumor efficiency [34], we investigated its efficacy on CSCs in vivo.

GNS561 has been previously shown to have potent antitumor activity in two HCC *in vivo* models including a diethylnitrosamine-induced cirrhotic rat model of HCC [14]. Here, we used the same model to study the impact of GNS561 on the frequency of CSCs *in vivo*. Rats with already developed HCC were either treated with vehicle, sorafenib (a standard of care of HCC) at 10 mg/kg or GNS561 at 15 mg/kg. The CD90^+^CD45^-^ phenotype was used to identify and quantify CSCs, since it has been reported to be associated with CSCs in liver tumor tissues and with circulating CSCs in blood samples [35–37]. In our rat model, no significant decrease was observed in the frequency of CSCs from the liver tumor tissue between the GNS561-treated group and the control group (Fig. 3A). However, the decrease in circulating CSCs became significant in the GNS561-treated group compared with the control group in the circulating CSCs (p = 0.0067, Fig. 3B). Notably, sorafenib, which is widely used to treat advanced HCC in humans, had no significant effect on the percentage of CSCs.

**Figure 3.**
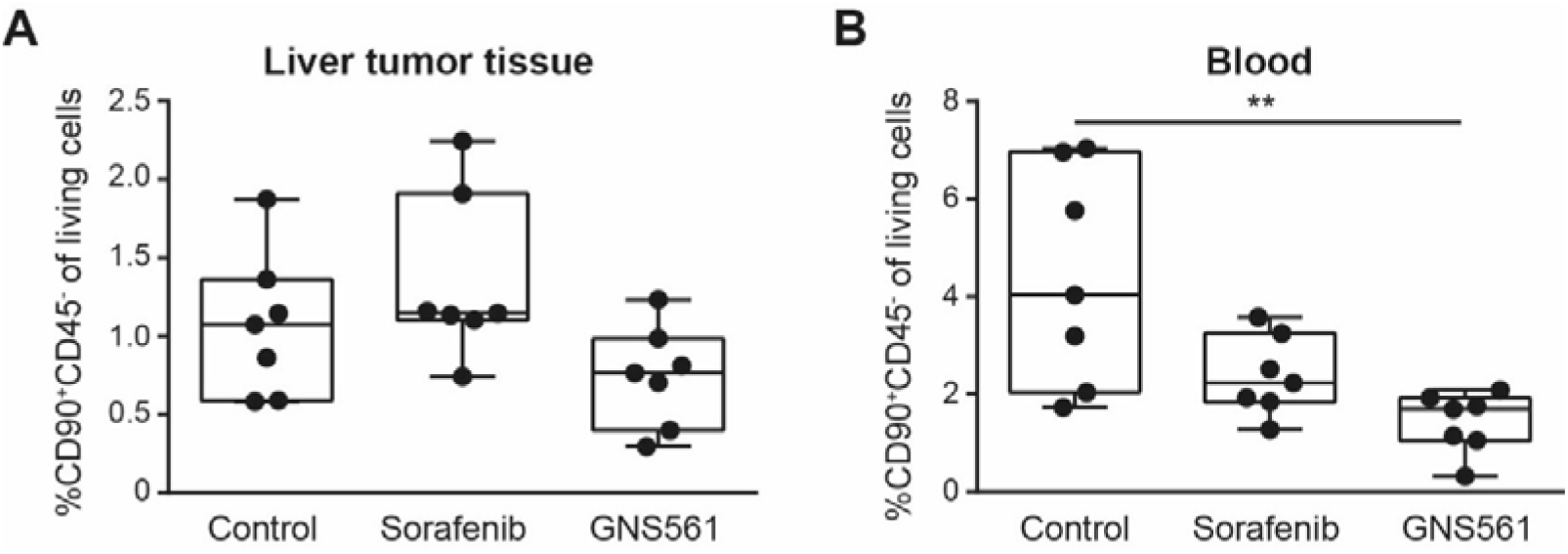
GNS561 decreases CSC frequency *in vivo*. Box and whisker representation (min to max) of the percentage of CSCs in liver tumor (A) and blood (B) samples in a diethylnitrosamine-induced cirrhotic rat model of HCC. Rats received vehicle (control), sorafenib at 10 mg/kg or GNS561 at 15 mg/kg. For comparison with the control non-treated group, the non-parametric Kruskal-Wallis test was performed. **, P < 0.01.

## Discussion

HCC and mCRC are refractory to most conventional chemotherapies. One of the reasons for the high mortality rate is the existence of a small population of CSCs with high tumorigenicity and resistance to drugs. Accordingly, many cancer therapies, while killing the bulk of tumor cells, may ultimately fail because they do not eliminate CSCs that reform heterogeneous tumors.

GNS561 is a small molecule that showed potent antitumor activity *in vitro* in a panel of human cancer cells, including HCC patient-derived cells, and *in vivo* in two HCC different models [14]. The *in vitro* studies performed in this study clearly showed that GNS561 displays potent dose response activity when assayed against a panel of human colon tumor cells isolated from liver metastases. In addition, GNS561 was efficient against subpopulation displaying ALDH^bright^ activity and produced a dramatic decrease in the ability of these cells to form tumorspheres. Similarly, in two HCC cell lines, GNS561 was equally effective in both populations (bulk and CSCs) and dramatically decreased the cell capacity to form spheres, even at low doses, unlike sorafenib, a standard of care. This latter observation was in agreement with the literature. Indeed, CSC resistance to sorafenib has been widely reported and has been cited as responsible for treatment failure and relapse [38,39].

Taken together, our results strongly suggest that GNS561 was efficient against both the whole tumor cell population and cell subpopulations displaying CSC features (ALDH^bright^ cells and tumorsphere initiating cells). These original results were confirmed *in vivo*, in a HCC-induced cirrhotic rat model, in which GNS561 showed antitumor effects and reduced the frequency of CSCs in liver tumor tissue and circulating CSCs. To go further, it would be informative to xenograft tumorsphere-derived CSCs or serial transplant tumors to determine GNS561 activity on the tumor growth.

The effectiveness of GNS561 to kill both tumor bulk cells and CSCs can be explained by its capacity to inhibit autophagy and lysosomal functions and to disrupt lysosomes [14]. In fact, once established, tumors require an uninterrupted nutritional supply for maintaining their proliferative needs in stressful conditions such as hypoxia, nutrient and growth factor starvation, and oxidative and metabolic stress [17]. In this context, autophagy has a protumoral activity by providing recycled bioenergetic substrates and consequently overcoming nutritional deficiency and favoring the survival of cancer cells [40]. Concerning the CSC population, autophagy was shown to be a major factor for the preservation of cell homeostasis and the maintenance of stemness properties [5]. Consistently, it was reported that metastatic cells and CSCs are vulnerable to lysosomal inhibition and disruption [41–44]. For example, inhibition of autophagic flux and lysosomal functions by salinomycin interferes with the maintenance and expansion of breast cancer stem-like/progenitor cells [45]. Moreover, a salinomycin derivative has been showed to be able to sequester iron in lysosomes ultimately resulting in cell death of CSCs [44,46]. Mefloquine showed activity on acute myeloid leukemia cells and progenitors by disrupting lysosomes (lysosomal membrane permeabilization and cytosolic release of cathepsins) [47].

Further research will be necessary to better understand the underlying mechanisms implicated in GNS561 activity against CSCs. It would be interesting to evaluate whether GNS561 treatment results from GNS561-induced inhibition of PPT1 and the mTOR pathway and lysosomal zinc sequestration in CSCs, as demonstrated in whole tumors [14]. In fact, knockout of PPT1 in tumor cells inhibited tumor growth, tumorsphere formation and tumorigenesis *in vivo* [48], suggesting that PPT1 may be implicated in CSC maintenance. In turn, the mTOR pathway has been shown to be one of the most important pathways involved in the development and progression of cancer and in the maintenance and hallmarks of CSCs [49], implying that inhibition of the mTOR pathway could be a good therapeutic strategy to eradicate CSCs. Moreover, as autophagy is an adaptive mechanism modulating the TME surrounding CSCs to support their stemness and cancer propagation [19], studying the effect of GNS561 on the TME would be of broad interest. Finally, it was recently reported that interfering with iron homeostasis using small molecules alters epigenetic plasticity of cells [50]. To evaluate if zinc could also be a regulator of epigenetic plasticity and if GNS561, by sequestrating lysosomal zinc, impairs this process would also be of particular interest.

In conclusion, GNS561 showed activity against not only the whole tumor population but also against subpopulations displaying CSC features *in vitro* and *in vivo*. Thus, a strategy that simultaneously exterminates the CSC subpopulation and tumor bulk by using autophagy inhibitors and lysosomal disruptors, such as GN561, is promising as a new direction to achieve cure and to prevent relapse of liver primary cancer and metastasis. GNS561 could also be used in combination with other drugs, in order to enhance the overall anticancer effect, as already described for hydroxychloroquine in several tumor types (reviewed in [3]). In particular, as classical chemo- or radiotherapy mainly fails to eliminate the CSCSs present in many tumor types, GNS561 could be useful to target these CSCs and then to reduce the risk of relapse in patients.

## Abbreviations

ALDH: Aldehyde dehydrogenase
CSCs: cancer stem cells
FBS: fetal bovine serum
HCC: hepatocellular carcinoma
IC_50_: half-maximal inhibitory concentration
mCRC: metastatic colorectal cancer
PPT1: palmitoyl-protein thioesterase 1
SEM: standard error mean
TME: tumor microenvironment.

## Acknowledgement

The authors are very grateful to Dr. Michel Prudhomme and Dr. Jean-François Bourgaux from CHU Carémeau, Nîmes, France, for the use of liver mCRC cells.

## Conflicts of Interest

SB, EB, FB, SM, JC and PH are employees of Genoscience Pharma. SB, FB and PH are shareholders of Genoscience Pharma. SB, FB, JC and PH are co-inventors of a pending patent. The other authors declare that they have no conflicts of interest to report.

## Funding

This work did not received external funding

## Notes

### Summary of Updates

To submit a new version of the manuscript

