## Supplementary Data for "GNS561, a New Autophagy Inhibitor Active against Cancer Stem Cells in Hepatocellular Carcinoma and Hepatic Metastasis from Colorectal Cancer"

### **Supplementary Materials and Methods**

*RNA extraction and real-time quantitative PCR (RT-qPCR)*

Total RNA was extracted using Tri-Reagent (SIGMA), according to the manufacturer’s instructions. About 20μg of total RNA was DNAse I‐digested (Roche), and then 1µg was reverse transcribed (RT) by using the Superscript IV First-Strand Synthesis System (Invitrogen) and random primers. cDNA. Amplification was carried out on 1/10 diluted cDNA with a LightCycler 96 instrument (Roche) using QuantiFast SYBR Green PCR Kit (Qiagen). Levels of gene expression were determined with GUSB gene as reference. The following primers were used: GUSB Forward 5’ CGT GGT TGG AGA GCT CAT TTG GAA 3’, Reverse 5’ ATT CCG CAG CAC TCT CGT CGG T 3’; NANOG Forward 5’ CAT CCC TGG TGG TAG GAA GA 3’, Reverse 5’ CCA ACA TCC TGA ACC TCA GC 3’; OCT4 Forward 5’ GTG AAG TGA GGG CTC CCA TA 3’, Reverse 5’ GAA GGA TGT GGT CCG AGT GT 3’; SOX2 Forward 5’ AAC CCC AAG ATG CAC AAC TC 3’, Reverse 5’ CGG GGC CGG TAT TTA TAA T 3’. Experiments were performed three times in duplicate.

*In vitro cell viability assay*

HCC and liver mCRC cell lines were seeded in 96-well plates at 10,000 cells/well. After 24 h, GNS561 or vehicle were added at appropriate concentrations. At the indicated time points, cell viability was assayed using CellTiter Cell Proliferation Assay (Promega) or CellTiter-Glo Luminescent Cell Viability Assay (Promega) following the manufacturer’s protocol.

For experiment comparing antitumor activity of GN561 on the whole population and CSC of HCC cell lines, spheres were dissociated by trypsination and seeded for cell viability assays as cells cultured in adherent conditions (10,000 cells/well). Cell viability determination was performed as described above.

Cell viability was expressed as a percentage of the values obtained from the negative control cells (vehicle treated cells). The half-maximal inhibitory concentration (IC_50_) was evaluated after 72 h treatment using a nonlinear regression curve in Prism 8.4.3 software. Each concentration was tested in triplicate. Mean IC_50_ was calculated as the average of three independent experiments.

*Aldehyde dehydrogenase activity assay*

The aldehyde dehydrogenase (ALDH) activity was determined using the ALDEFLUOR™ assay (Stem Cell Technologies) according to the manufacturer’s instructions. ALDH^bright^ cells were identified by comparing the same sample with and without the ALDH inhibitor diethylaminobenzaldehyde (DEAB). Gating of ALDH activity was set at 0.1% in presence of DEAB. Cells were analyzed using MacsQuant (Miltenyi Biotec) and data analyzed using Flowing softwares (CyFlo Ltd). CPP36 cells were not used in these analyses because of their high cellular autofluorescence profile*.*

*Tumorsphere formation assay*

Evaluation of tumorsphere formation ability of HCC cell lines was determined after seeding cells (12 500 cells / 1mL) in sphere medium (complete cell culture medium without FBS and supplemented with B27 (Life Technologies), 20 ng/mL EGF (R&D Systems), 20 ng/mL bFGF (StemCell Technologies) and 4 µg/mL heparin (Sigma) and distributed into ultra-low adherence 24 wells cell culture plates (Corning). At the time of seeding, GNS561, sorafenib (Santa Cruz Biotechnology) or vehicle were added to the media. Sphere size exceeding 50 µm were counted after 96 h by using the Icy software (Pasteur Institute, France; http://icy.bioimageanalysis.org/) and represented at number of spheres per well.

Evaluation of tumorsphere formation ability of liver mCRC cell lines was determined after plating 500 cells / 200 µL well in M11 medium in P96 wells in ultra-low attachment plates (Corning). M11 is DMEM/F12 (1:1) medium (Gibco), supplemented with N2 (Invitrogen), 3 mM glutamine (Gibco), 0.6% glucose (Sigma-Aldrich), 4 µg/ml insulin (Sigma-Aldrich), hBasic-FGF 10 ng/ml (R&D Systems), and hEGF 20 ng/ml (R&D Systems). GNS561 (0.3 or 3 µM) was added 24 h after cell plating in M11 medium. Sphere size exceeding 50 µm were counted after 10 days (NIS-Elements, Nikon) and represented at number of spheres per image field. CPP45 cells were not used in these analyses because of their inability to from tumorospheres.

*Secondary tumorsphere formation*

Primary spheres from Hep3B were isolated by filtration through a 70 µM cell strain (Falcon), trypsinized and seeded into ultra-low adherence 24 wells cell culture plates, as described above.

*Diethylnitrosamine-induced immunocompetent rat model of HCC*

Thirty 6-week-old Fischer 344 male rats (Charles River Laboratories) were housed in the animal facility of Plateforme de Haute Technologie Animale (Jean Roget, University of Grenoble-Alpes). Rats were kept in individually ventilated cage systems at constant temperature and humidity with 2-3 animals in each cage having free access to food (standard diet) and water during the entire study period. Rats were treated weekly with intra-peritoneal injections of 50 mg/kg of diethylnitrosanime, which were diluted in olive oil in order to obtain a fully developed HCC on a cirrhotic liver after 14 weeks [1]. Rats were randomized in 4 different groups and treated during six weeks by i) sorafenib (n=8), ii) GNS561 (n=8), iii) vehicle (control, n=8). Treatments were administered by oral gavage for a period of six weeks. The nutritional state was monitored by daily weighing of rats and protein-rich nutrition was added to the standard food in every cage where a loss of weight was observed. The food intake per cage was monitored during the last 6 weeks of the experiment. Food was withheld for 3-4 h before the animals were sacrificed. Imaging study was conducted on a 4.7 Tesla MR Imaging system (BioSpec 47/40 USR, Bruker Corporation. Cells were recovered from liver tumor tissue by mechanical disruption and whole blood samples were used in case of blood analyses. Cells were immunostained for flow cytometric analysis without any stimulation. Following anti-rat antibodies were used: CD45 (APC/Cy7 - BioLegend) and CD90 (BV711 - BioLegend). Nonviable cells were stained by Zombie UV™ Fixable Viability Kit (BioLegend) and excluded from the analysis. BD CompBead particles were used for compensation and isotype-matched antibodies as negative control. Data were acquired on BD-LSRII flow cytometer (BD Biosciences), collected with BD FACSDiva 6.3.1 software and analyzed using FCS Express 6 PLUS software.

### **Figure S1. Hep3B tumorspheres overexpress stemness-associated genes compared to the total adherent population.**

Hep3B cells were dissociated and seeded either in adherent or non-adherent culture conditions. After 96h, cells were harvested. RT-qPCR was performed on total RNA (see materials and methods). The figure represents the mean of three independent experiments. Man-Whitney statistical test was used to compare tumorspheres and adherent cells. *, p < 0.05.

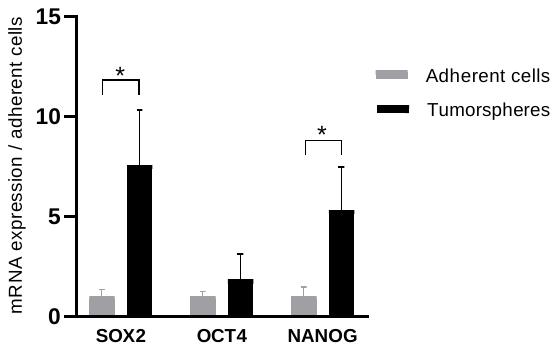

### **Figure S2. GNS561 is efficient against subpopulation displaying CSC features.**

Antitumor activity of GNS561 on whole tumor (circles) or CSC-enriched populations (squares) in Hep3B after 6 (A), 8 (B) and 24h (C) of treatment with GNS561. The curves represent the mean of two independent experiments in triplicate.

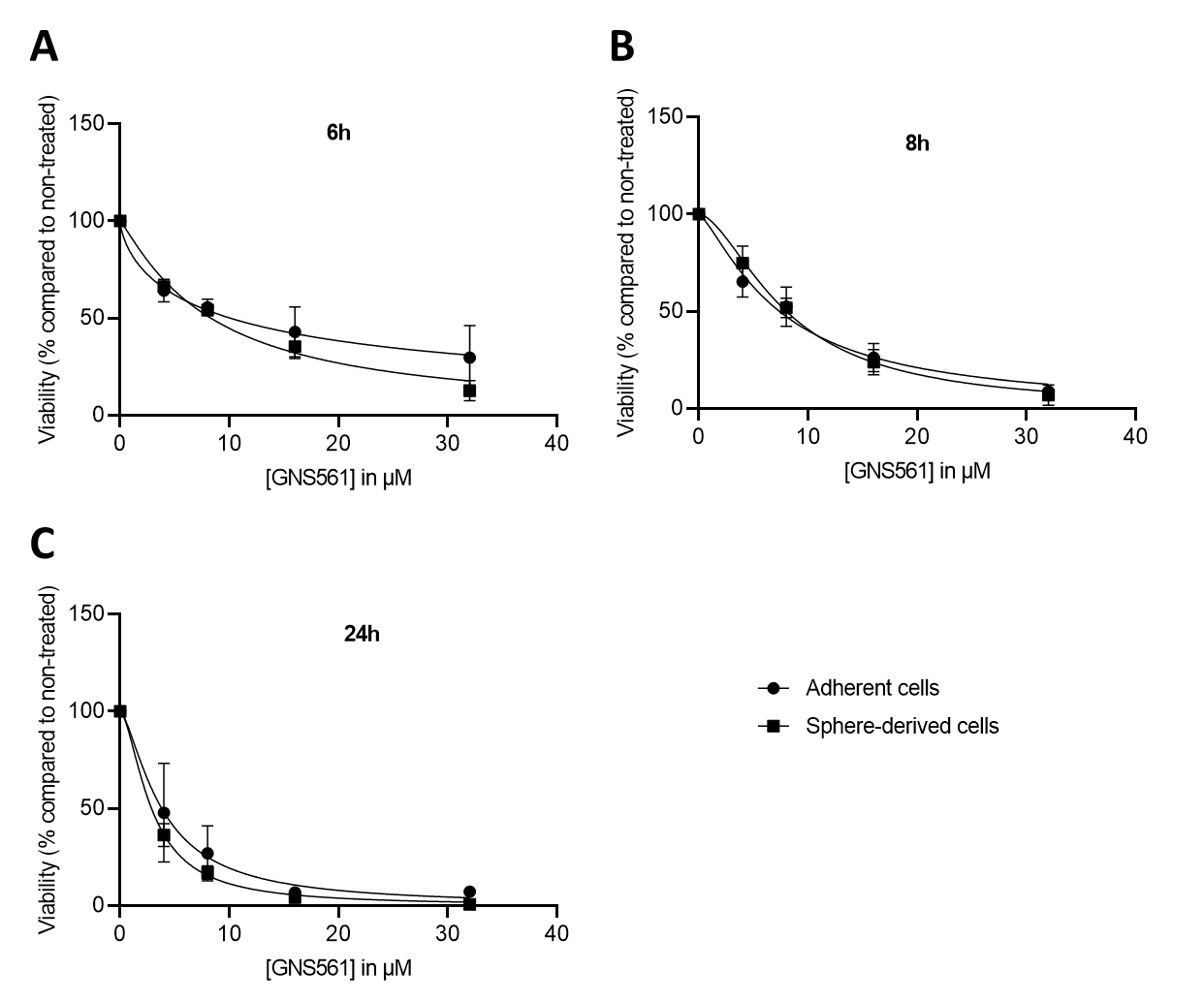
**Figure S3. GNS561 but not sorafenib alters the ability of Hep3B to form secondary spheres.**

Hep3B primary spheres were dissociated and seeded in non-adherent culture conditions in presence or not of GNS561 (A) and sorafenib (B). Phase-contrast microscopy examination of cells after four days of culture is shown. Pictures are representative of two independent experiments in duplicate. Scale bar, 50µM.

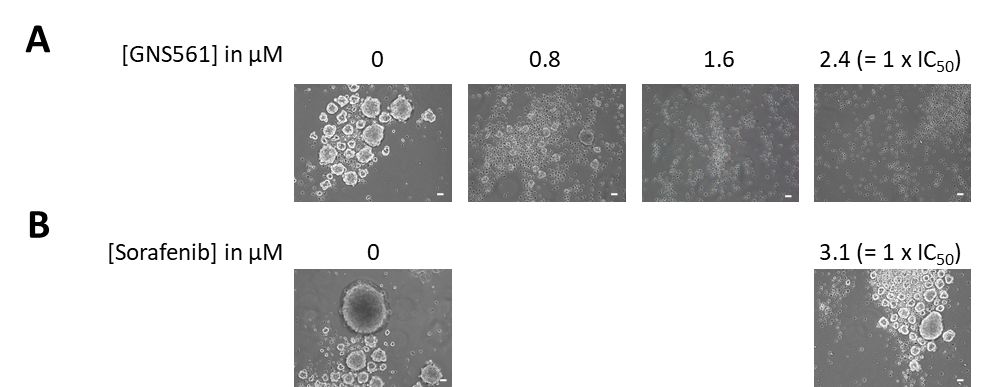

### **Figure S4. Sorafenib does not abolish the tumorsphere formation ability of Hep3B.**

Hep3B adherent cells were dissociated and seeded in non-adherent culture conditions in presence or not of 1 x IC_50_ sorafenib. Phase-contrast microscopy examination of cells after four days of culture is shown. Pictures are representative of three independent experiments in duplicate. Two fields are presented for each condition. Scale bar, 50 µM.

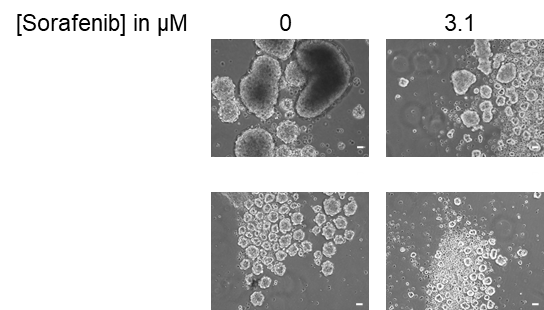
